# Champagne: Whole-genome phylogenomic character matrix method places Myomorpha basal in Rodentia

**DOI:** 10.1101/803957

**Authors:** James Kusik Schull, Yatish Turakhia, William J. Dally, Gill Bejerano

## Abstract

We present Champagne, a whole-genome method for generating <u>cha</u>racter <u>m</u>atrices for <u>p</u>hylogenomic <u>a</u>nalysis using large <u>g</u>e<u>n</u>omic <u>e</u>vents that, by rigorously picking orthologous genes and locating large, virtually homoplasy-free insertion and deletion events, delivers a character matrix that outperforms existing morphological and nucleotide-based matrices on both established phylogenies, and difficult-to-resolve nodes in the mammalian tree. Champagne harbors distinct theoretical advantages, and can easily be run on any clade of related species, of the many currently being sequenced. Champagne considerably improves the retention index in the parsimony analysis of a number of widely established topologies, observes incomplete lineage sorting (ILS) at the root of Paenungulata, finds little evidence for human-chimp-gorilla ILS, and most surprisingly, offers convincing evidence for a reconsideration of squirrel’s position in the rodent tree.

## Introduction

The “phylogenomics” approach^1^ promises to resolve the branching patterns in the tree of life with the enormous statistical power of genome-scale data. Many recent phylogenomic studies have confirmed topology inferences of previous studies that mostly relied on morphological features^2^, while others have led to new revisions to our current understanding of the tree of life^3–5^.

Yet, despite the proliferation of high-quality whole genome assemblies, many topologies in the mammalian tree remain hotly contested in phylogenomic studies^6–10^. Phylogenomic studies reconstruct phylogenetic trees from a character matrix composed of molecular signals, such as DNA or protein alignments, which suffer from a number of stochastic and systematic biases that sometimes result in species tree incongruence^11^. Stochastic biases can arise from biological mechanisms, such as incomplete lineage sorting (ILS)^12, 13^, hybridization^14^ and horizontal gene transfer^15^, as well as algorithmic shortcomings, such as alignment issues and incorrect orthology mapping. Systematic biases may result from homoplasy — increased rate of parallel or convergent mutations which might be a result of mutation rate-heterogenity^16, 17^ or similar selective pressures^18^. As noted by Jeffroy et al.^11^, contrary to stochastic bias, systematic bias can be reduced not by adding more signal to the character matrix, but by reducing the level of non-phylogenetic signal in the character matrix. A similar argument has been made by Philippe et al.^19^. While there is a rich body of recent literature focused on statistical techniques to reduce stochastic bias in phylogenetic analysis, particularly from ILS^20, 21^ and hybridization^22, 23^, very little work has been done to generate a homoplasy-free character matrix for phylogenetic analysis from entire genomes.

In this paper, we present Champagne — a method for generating <u>cha</u>racter <u>m</u>atrices for <u>p</u>hylogenetic <u>a</u>nalysis using large <u>g</u>e<u>n</u>omic <u>e</u>vents. Champagne builds a character matrix using large (*>*75bp) shared insertions and deletions (indels, in short) within the introns of orthologous genes among the species of interest using gene annotations in a known outgroup species. This has two major advantages over prior techniques. First, by using large shared insertions and deletions, which are extremely unlikely to occur independently, Champagne largely eliminates homoplasy that is prevalent in single nucleotide (or amino acid) level DNA (or protein) alignments, where parallel and convergent mutations occur frequently. Second, while some prior work has focused on large shared genomic regions for inferring phylogeny with promising results, such as using SINEs^24^, ultraconserved elements^25^ and retroposons^26^, their techniques are typically manually curated for specific regions in the genome and discover only a handful of informative sites, which might be less significant statistically or have sampling biases. Champagne is fully-automated, works on raw genome sequences of target species, and typically discovers hundreds to tens of thousands of informative sites, including many in the non-coding portions of the genome. Traditionally, it has been challenging to establish orthology in non-coding portions of the genome. To address this issue, Champagne uses a strict algorithm for mapping each reference gene to at most a single orthoglous query locus and using pairwise alignment to further restrict the search to intragenic regions (containing both exons and introns).

When applied to mammalian genomes, Champagne improves confidence in inferring well-established topologies, producing character matrices with significantly lower homoplasy than the matrices presented in recent morphological and nuclear sequence-based phylogenetic studies. It discovers surprising but compelling evidence to position Myomorpha basal to Sciuridae and Hystricomorpha and reaffirms the high prevalence of ILS and similar effects related to rapid speciation in Paenungulata, even in considering large genomic events.

## Results

### Champagne identifies numerous shared large indels between species to produce its “homoplasy-free” character matrix

Champagne is a fully-automated, multi-stage computational pipeline that produces a set of phylogenetically informative evidence of large (*>*75bp) shared indels in the NEXUS format^27^, thus permitting the subsequent use of any chosen topology inference algorithm^28–32^. Champagne requires a single known outgroup with an annotated gene set and raw genome assemblies for the ingroup (also referred to as query) species. Figure 1 illustrates the Champagne algorithm. The pipeline consists of a series of discrete stages. Once a set of species (including an outgroup) has been selected, Champagne constructs new or uses available alignment chains (referring to the UCSC pairwise alignment *chains*^33^, see Methods) for all outgroup-query pairs, using those chains to map each outgroup gene to at most one orthologous chain in each query species (see Figure 1, step 1, and Methods for details). Ambiguous mappings are discarded. Next, for each outgroup gene that maps uniquely to more than one query species, Champagne scans the orthologous query regions corresponding to the outgroup intragenic region (exons and introns, where orthology is established with high-confidence), moving through the outgroup-query chains simultaneously and identifying large one-sided gaps in the chains (implying either an insertion in query or a deletion in outgroup, or vice versa). Upon finding this gap, Champagne determines whether this site could be phylogenetically informative i.e. at least two species could be found containing the sequence corresponding to the one-sided gap with high sequence similarity and at least two species could be found with an absence of that sequence (see Methods and Supplementary Figure 1 for details). By the parsimony argument, we assume that the ancestral (common to ingroup and outgroup species) state (presence or absence of that sequence) is the same as the state of outgroup species (Figure 1, step 2): for this to be false, the indel corresponding to that sequence would have had to independently occur at least twice, once in the outgroup and once in the ingroup species sharing the outgroup state. Since it is extremely unlikely that two large indels of roughly the same sequence would independently occur at the same locus, this parsimony assumption is relatively safe to make. By the end of this step, for each informative site, all ingroup and outgroup species are assigned a character state of ‘+’, ‘-’ or ‘?’, depending on whether the specific indel sequence of interest is present, absent or cannot be confidently determined in that query species, respectively. Each informative site is classified as a shared insertion or deletion between the query species differing from the ancestral and are written to an output NEXUS file (Figure 1, step 3). Finally, Champagne finds the most parsimonious topology from the NEXUS file using PAUP*’s maximum parsimony algorithm^28^, although alternative topology inference tools^29–32^ could also be used at this step (Figure 1, step 4).

**Figure 1.**
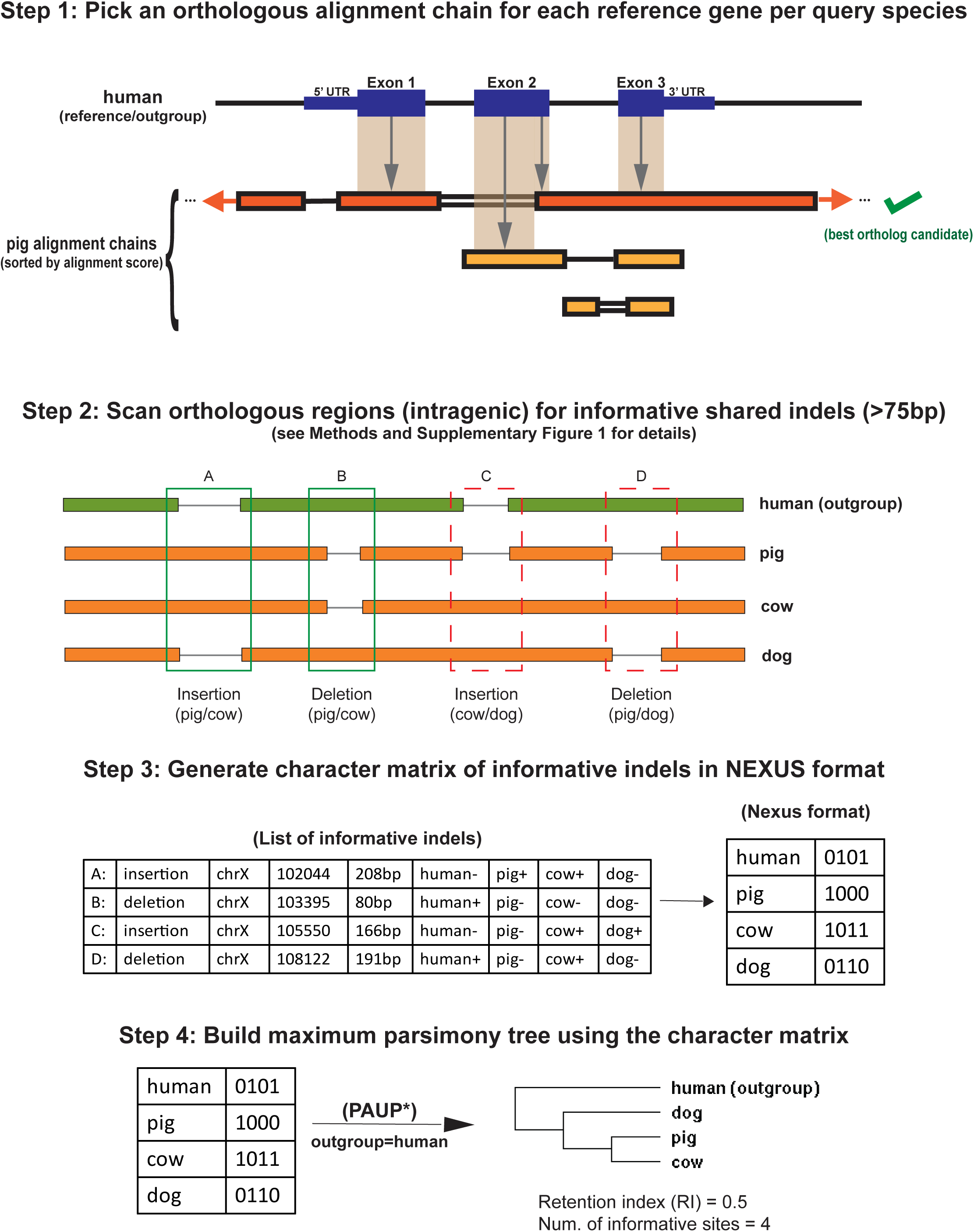
An overview of the Champagne approach for speciation topology inference. In step 1, we use pairwise alignment chains between the outgroup (also used as reference) and each ingroup species (used as query) to assign at most one orthologous chain with high-confidence for each reference gene. The figure illustrates this procedure for a single outgroup-ingroup pair (human-pig) and a single reference gene. Each coding base-pair in the gene is assigned to the highest-scoring chain overlapping with the gene. If the highest-scoring overlapping chain also has the most base-pairs assigned, it is chosen as the best ortholog candidate (as shown). If *gene-in-synteny* and *1-to-1 mapping* criteria are also satisfied (see Methods), the best candidate chain is assigned as gene ortholog. In all remaining cases, no assignment is made. In step 2, intragenic orthologous regions in all query species are scanned for each reference gene in search of phylogenetically informative, shared indels within ingroup (see Methods and Supplementary Figure 1 for details). In our illustration, four informative indels (labelled *A*, *B*, *C* and *D*) are found. In step 3, the informative indels are printed to a NEXUS file, which is used in step 4, to infer the most parsimonious species tree, here ((pig, cow), dog), using PAUP*^28^. Indels *A* and *B* in step 2 provide supporting evidence for ((pig, cow), dog), as only pig and cow share both indels. The other two indels, *C* and *D*, support ((cow, dog), pig) and ((pig, dog), cow) trees as most parsimonious, respectively. The low retention index (0.5 of maximum 1) of these four site examples reflects the relatively large fraction of non-supporting evidence in this topology assignment.

Figure 2 further illustrates a 14Mbp region in the human (outgroup) genome with real indel events annotated by Champagne for the species set *{*pig, cow, dog*}*. Even in this short segment, Champagne finds more indels shared by pig and cow, not observed in dog and human (outgroup), which support the most parsimonious topology (in Newick format): ((pig, cow), dog).

**Figure 2.**
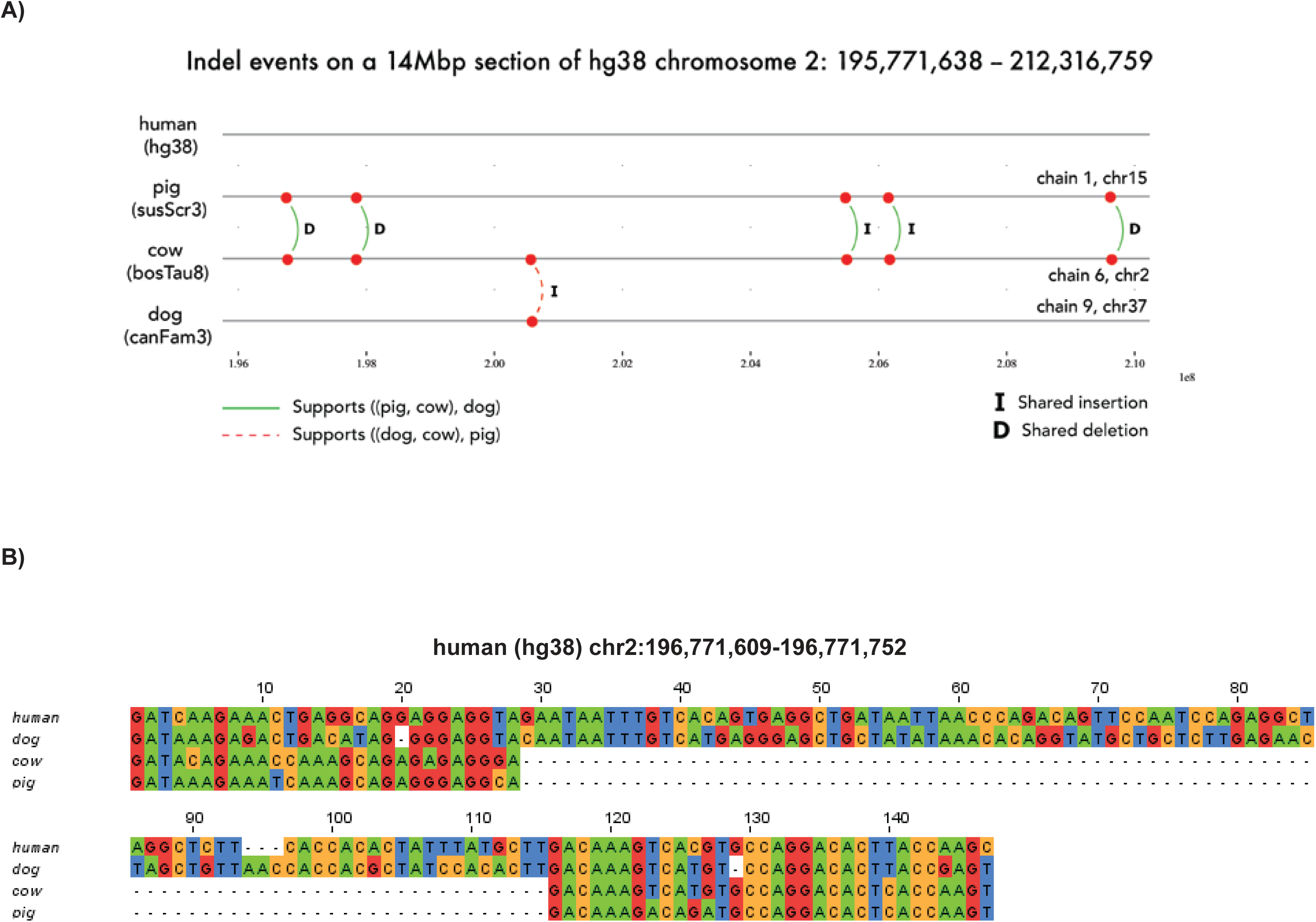
A multiple-species-alignment showing indels identified by Champagne in the pig, cow, and dog genomes, using human as reference species. **A)** An illustration of the real pig, cow, and dog chains that align with a 14Mbp section of the human chromosome 2. Indels identified by Champagne in this section of the reference genome are shown: “I” indicates shared insertions, and “D” indicates shared deletions. On this stretch, we find 6 indels that are shared by pig and cow, supporting the most parsimonious topology ((pig, cow), dog), and only 1 (shown with a dashed arc) that is shared by dog and cow, possibly due to ILS. **B)** A multiple sequence alignment of an 81bp deletion shared by pig and cow, but not dog (leftmost deletion in panel A).

Champagne does not suffer from a considerable long branch attraction^34^ (a phenomenon common in single nucleotide and amino acid space, whereby one or more species with a high mutation rate introduce a systematic error in phylogenetic analyses due to frequent convergent and reversal mutations), as large indel events in Champagne matrices are unlikely to occur independently or be reversed. For this reason, the maximum parsimony algorithm is indeed suitable for Champagne, particularly in the absence of an explicit evolutionary model for large indels, an equivalent of K80^35^ or F81^30^ used in nucleotide substitution, that is necessary for Maximum Likelihood or Bayesian inference approaches. We use the retention index (RI) yielded by PAUP* from the most parsimonious topology as an overall measure of the goodness-of-fit of Champagne’s character matrix to the optimal phylogeny. The retention index, first proposed by Farris^36^ in 1989, expresses the degree of synapomorphy (characters shared by descendants of a common ancestor) in a character matrix; it has been interpreted as a metric for assessing the degree to which a character matrix fits a given topology, and has been widely used since to support phylogenies^37^. Since the retention index reflects a normalized value (between 0 and 1) corresponding to the number of state changes required along the branches of a given phylogenetic tree to fit the character states along the tree’s leaves while also considering the theoretical best and worst case for the same character states, it can also be interpreted as a measure of apparent homoplasy (with higher values implying *lower* homoplasy) in a dataset. While RI is a powerful metric to quantify the aggregate homoplasy of a character matrix to a phylogeny, it may not clearly reflect the goodness-of-fit for specific bifurcations internal to the tree, especially when more than three species are used. To overcome this, in this paper, we identify informative sites in the NEXUS file that support each bifurcation internal to the parsimonious topology, and for contentious bifurcations, use a similar method to identify informative sites, if any, that support alternative bifurcations (see Methods). The more supporting evidence found for a particular bifurcation relative to its alternatives, the more confidence can be attributed to it.

### Champagne significantly improves the retention index (RI) in parsimony analyses of established topologies over morphology- and short sequence-based matrices

To evaluate Champagne’s performance in producing evidence that yields the correct topology, we started with the simplest case: sets of three species. We chose six species sets for which the topologies are broadly accepted. A number of previous papers, building topologies on the basis of molecular and morphological datasets, have established the correct phylogenies for these species sets (presented in Newick format) to be: ((mouse, rat), guinea-pig); ((dog, cat), pig); ((dolphin, cow), horse); ((pig, cow), dog); ((megabat, microbat), dog); and ((human, mouse), dog)^2, 25, 38–41^. We summarize these phylogenies, including the outgroups used by Champagne, in Table 1. We note that of the six species sets we consider, the correct topology for human, mouse, dog is perhaps the most debated — some papers^9, 42^ have proposed the alternate topology of ((human, dog), mouse), though the broader consensus is still in favor of ((human, mouse), dog). We compare the indel-based character matrices produced by Champagne with a morphological character matrix presented by O’Leary et al.^43^ and a nuclear DNA based character matrix presented by Song et al.^39^. Because of the limited set of taxa available in O’Leary et al. matrix, we could not compare retention indices across all phylogenies. We found that on all six sets, Champagne, as well as Song et al. matrices, produced the same topologies with maximum parsimony, which also matched with the broadly accepted topologies in previous studies. O’Leary matrices also predicted the same topologies on two out of three topologies we could evaluate, but incorrectly predicted the ((dolphin, cow), horse) topology as ((cow, horse), dolphin). The matrices differed in their retention index (RI) scores and the number of informative sites (Table 1). In general, nucleotide substitution based matrices from Song et al. had far more characters than the morphological matrices of O’Leary et al. or the Champagne matrices, which are based on rare, large indel events. Despite this, the character matrices produced by Champagne significantly outperform both Song et al.’s and O’Leary et al.’s matrices, producing a retention index close to the maximum possible value of 1 in almost all cases (Table 1). This is because large genomic events that Champagne considers rarely occur twice independently, which is neither true of morphological characters nor base-pair substitutions.

**Table 1.** A comparison of the retention indices (RI, ranging between 0 and 1) and the number of informative sites of the maximum-parsimony trees generated using a single nuceleotide-based character matrix by Song et al^39^, a morphological character matrix by O’Leary et al.^43^ and our indel-based character matrix of Champagne. Champagne’s high to near-maximal RI across all six queries shows how resilient large indel based inference is to homoplasious events, exemplifying the desirable reduction of non-phylogenetic signal in the character matrix.

|  | Outgroup | Retention Index (RI) |  |  | Number of informative sites |  |  |
| --- | --- | --- | --- | --- | --- | --- | --- |
|  |  | O'Leary et al. | Song et al. | Champagne | O'Leary et al.<br>(Morphological traits) | Song et al.<br>(Single bases) | Champagne<br>(Large shared indels) |
| ((mouse, rat), guinea-pig) | Human | N/A | 0.84 | **0.997** | N/A | 55,922 | 295 |
| ((dog, cat), pig) | Human | N/A | 0.598 | **0.993** | N/A | 19,872 | 998 |
| ((dolphin, cow), horse) | Human | 0.445<br>(incorrect) | 0.657 | **0.99** | 155 | 29,708 | 306 |
| ((pig, cow), dog) | Human | 0.469 | 0.554 | **0.989** | 350 | 26,331 | 359 |
| ((megabat, microbat), dog) | Human | 0.581 | 0.481 | **0.933** | 296 | 22,942 | 45 |
| ((human, mouse), dog) | Elephant | N/A | 0.358 | **0.765** | N/A | 28,648 | 17 |

N/A: not available

### Champagne shows considerable effect of ILS in cross-species structural variation in species that underwent rapid radiation

Despite a proliferation of genomic data, many topologies in particular remain unresolved to this day hindered by rapid speciation and a corresponding prevalence of incomplete lineage sorting (ILS)^8^ (see Supplementary Figure 2). A classic example is the confounding branching pattern within Paenungulata (containing the clades Hyracoidea (hyraxes), Sirenia (manatees, dugongs, sea cows) and Proboscidea (elephants)). Several past papers have proposed contradictory tree topologies for Paenungulata, with some arguing that Hyracoidea is basal to Sirenia and Proboscidea^26, 44–46^, and others arguing that Proboscidea is basal^38, 47^. Of these, only Nishihara et al.^26^ studied this phylogeny using structural genomic changes involving retroposons but found only one informative site supporting Hyracoidea in the basal position. We sought to explore whether ILS effects resulting from the rapid radiation within Paenungulata could be observed on structural genomic changes using Champagne. We selected a compact set of species to represent each tree, and used Champagne to produce corresponding evidence matrices. For Paenungulata, we consider the minimal set: *{*elephant, manatee, rock hyrax*}*, with human as outgroup. The maximum parsimonious tree produced by Champagne supports the basal placement of Hyracoidea relative to Proboscidea and Sirenia (Figure 3). In particular, Champagne finds 422 indels supporting the topology: ((elephant, manatee), hyrax) (Figure 3A,B,C). In contrast, Champagne finds only 54 indels supporting the topology: ((hyrax, manatee), elephant), and 242 indels supporting the topology: ((elephant, hyrax), manatee). The unmistakable prevalence of ILS, evidenced by the relatively high proportion of indels identified that support the other possible hypotheses (Figure 3A,D), supports prior arguments concerning the difficulties of phylogenomic analysis at ILS-prone soft polytomous nodes and suggests that confident resolution of this topology will remain difficult for any amount of data or approach, as the conflicting signal is likely phylogenetic. However, while support for alternate hypotheses certainly reflects the heavy influence of ILS, the number of indels identified that support the most parsimonious tree is also considerably more compelling than the support for the less parsimonious trees. In this respect, we believe that Champagne produces a character matrix that relatively confidently supports the placement of Hyracoidea basal to Proboscidea and Sirenia. To our knowledge, Champagne also provides the first character matrix to observe prevalence of ILS on large cross-species structural variations.

**Figure 3.**
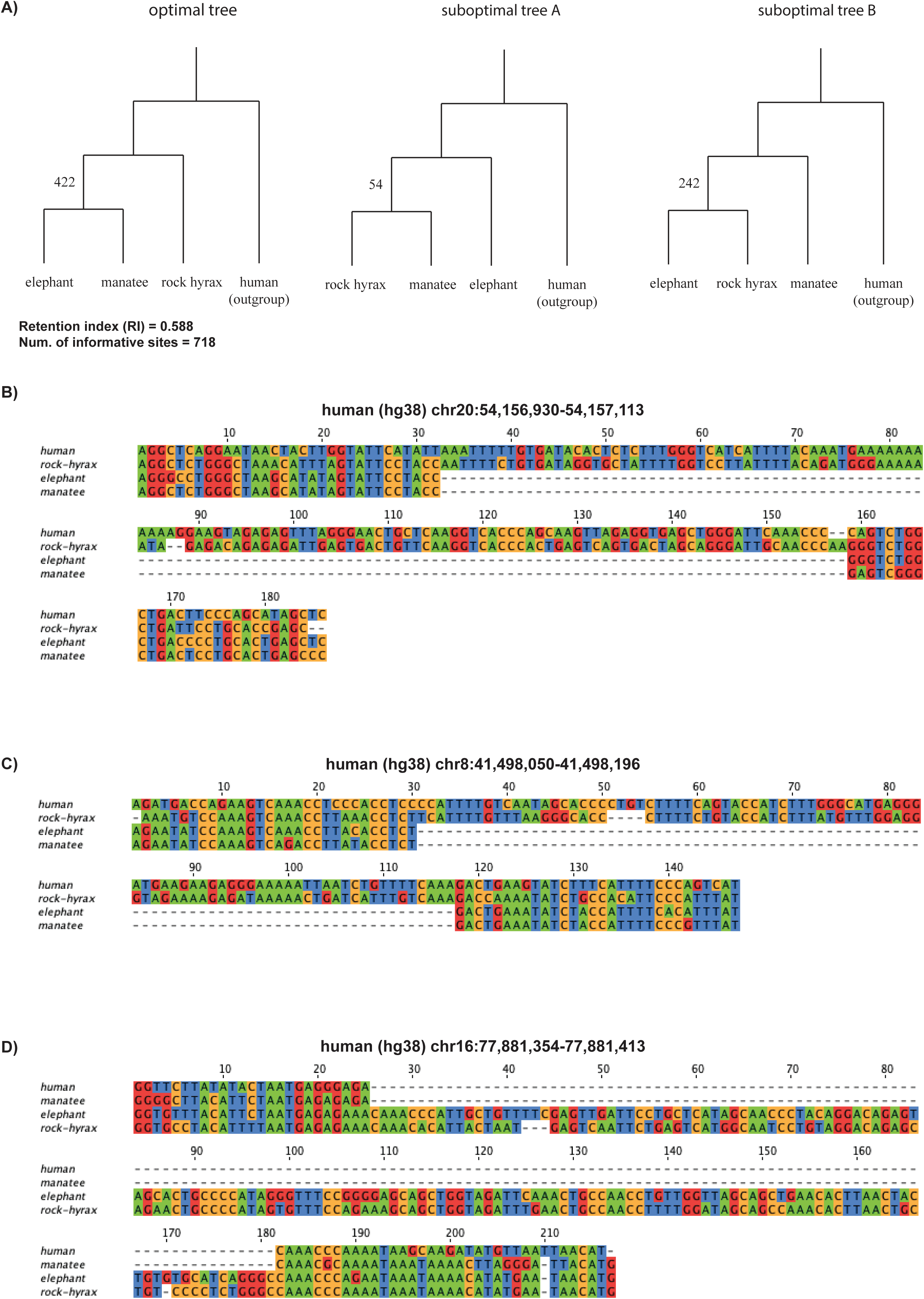
Champagne supports Hyracoidea as basal in the ILS heavy Paenungulata tree. **A)** The maximum parsimony tree generated by PAUP* using Champagne’s character matrix for Paenungulata (rock hyrax, (elephant, manatee)), as well as the other two less parsimonious alternatives. The high number of Champagne supporting indels per topology (and a moderate retention index) likely reflect incomplete lineage sorting (ILS) at the root of this subtree. **B)** A multiple sequence alignment for a 124bp deletion shared by elephant and manatee, one of 422 that supports our maximum parsimony topology. **C)** A multiple sequence alignment for an 87bp deletion shared by elephant and manatee that also supports our maximum parsimony topology. **D)** A multiple sequence alignment for a 152bp insertion shared by elephant and rock hyrax, supporting the topology ((elephant, rock hyrax), manatee), or strong ILS at the Paenungulata root.

### Champagne scales well to a larger number of species

By designing the indel-search algorithm to only involve outgroup-query chains, Champagne requires only linear time and *N* computationally-expensive chains to be produced for a phylogeny containing *N* species. For primates, we build a larger Champagne matrix containing the 9 primate species: *{*human, chimpanzee, gorilla, orangutan, macaque, marmoset, tarsier, galago, mouse lemur*}*, with mouse as outgroup (Figure 4). The maximum parsimony topology yielded by Champagne’s character matrix for these primates matches the topology inferred in a number of previous papers^25, 39, 41^ with a large number of supporting cases for most bifurcations (Figure 4A,C). Most importantly, 89 indels support grouping human and chimpanzee together before grouping either of them with gorilla or some other ingroup species, while 0 indels support the placement of chimpanzee basal to human and gorilla ((Figure 4B).

**Figure 4.**
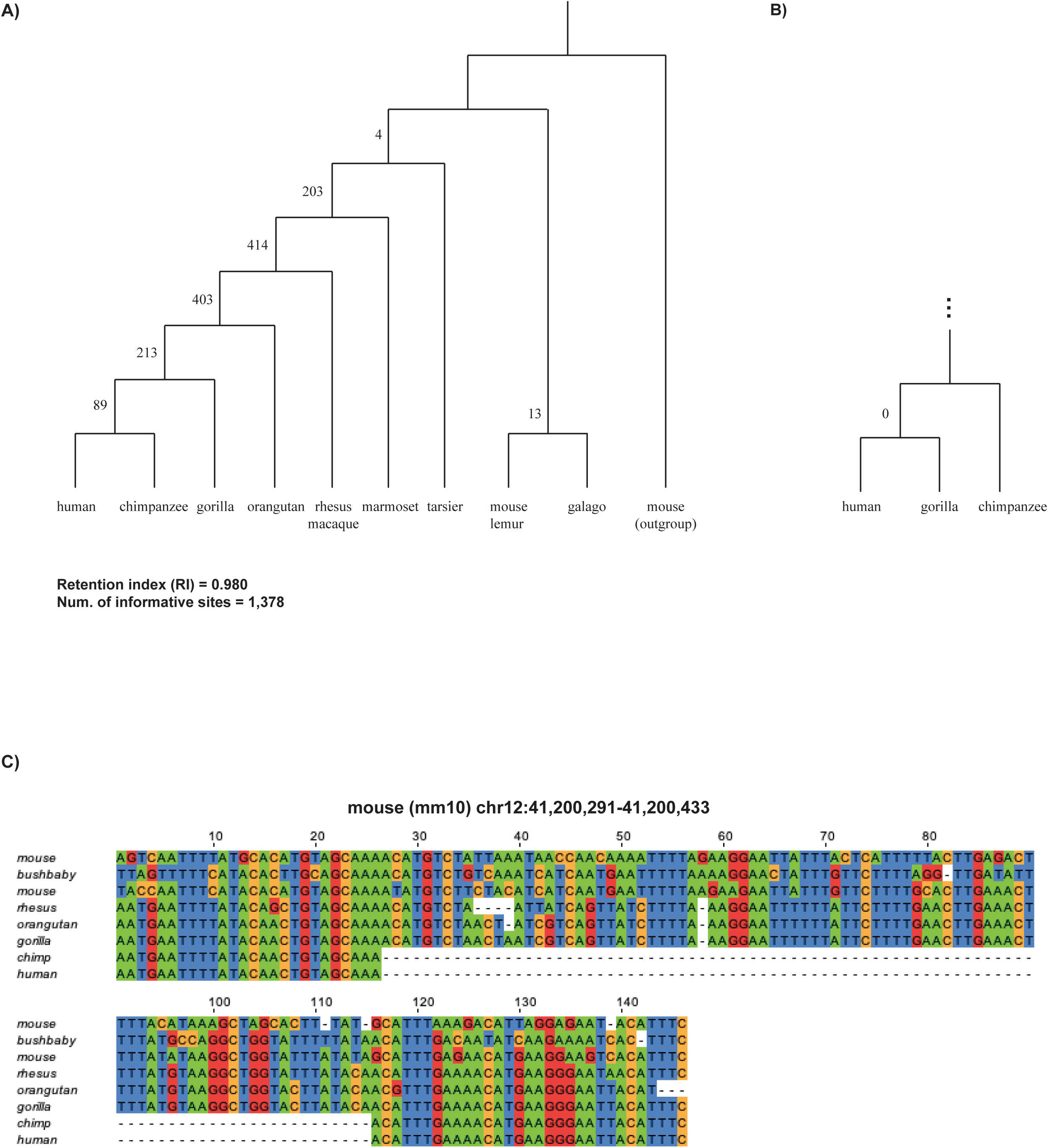
Champagne correctly reconstructs primate phylogeny, finding little evidence for human-chimp-gorilla ILS. **A)** At each node in the tree, we depict the number of indels identified by Champagne that support the corresponding clade. **B)** In particular, Champagne finds 0 indels supporting chimpanzee as an outgroup to human and gorilla, putting in question extensive ILS at this node. **C)** A multiple sequence alignment for an 87bp deletion shared uniquely by human and chimpanzee.

### Champagne provides surprising evidence to support Myomorpha basal to Hystricomorpha and Sciuridae

The relationship between Myomorpha (the clade that includes mouse and rat), Hystricomorpha (the clade that includes guinea-pig), and Sciuridae (the family containing squirrels) has also been debated in prior literature, with published phylogenies alternately presenting Myomorpha in the basal position^42^, Hystricomorpha in the basal position^25^, and Sciuridae in the basal position^6, 24^. To our understanding, recent consensus favoring Sciuridae in the basal position has emerged. Using the genomes of the species *{*mouse, rat*}* for Myomorpha, *{*naked mole rat, guinea-pig*}* for Hystricomorpha and *{*squirrel, marmot*}* for Sciuridae, we sought to explore this disputed topology using Champagne. To our surprise, we found significant evidence to place Myomorpha in the basal position, contrary to the latter recent studies, discovering 70 indels that support our phylogeny (Figure 5A,B,C). In contrast, we find 8 indels supporting the placement of Hystricomorpha in the basal position, and only 3 indels supporting the placement of Sciuridae in the basal position. Clearly, there is some ILS on the disputed node (Figure 5A,D), but the weight of evidence supporting the placement of Myomorpha in the basal position provided by Champagne is far more significant, and we believe it is compelling enough to revisit the current consensus on this phylogeny.

**Figure 5.**
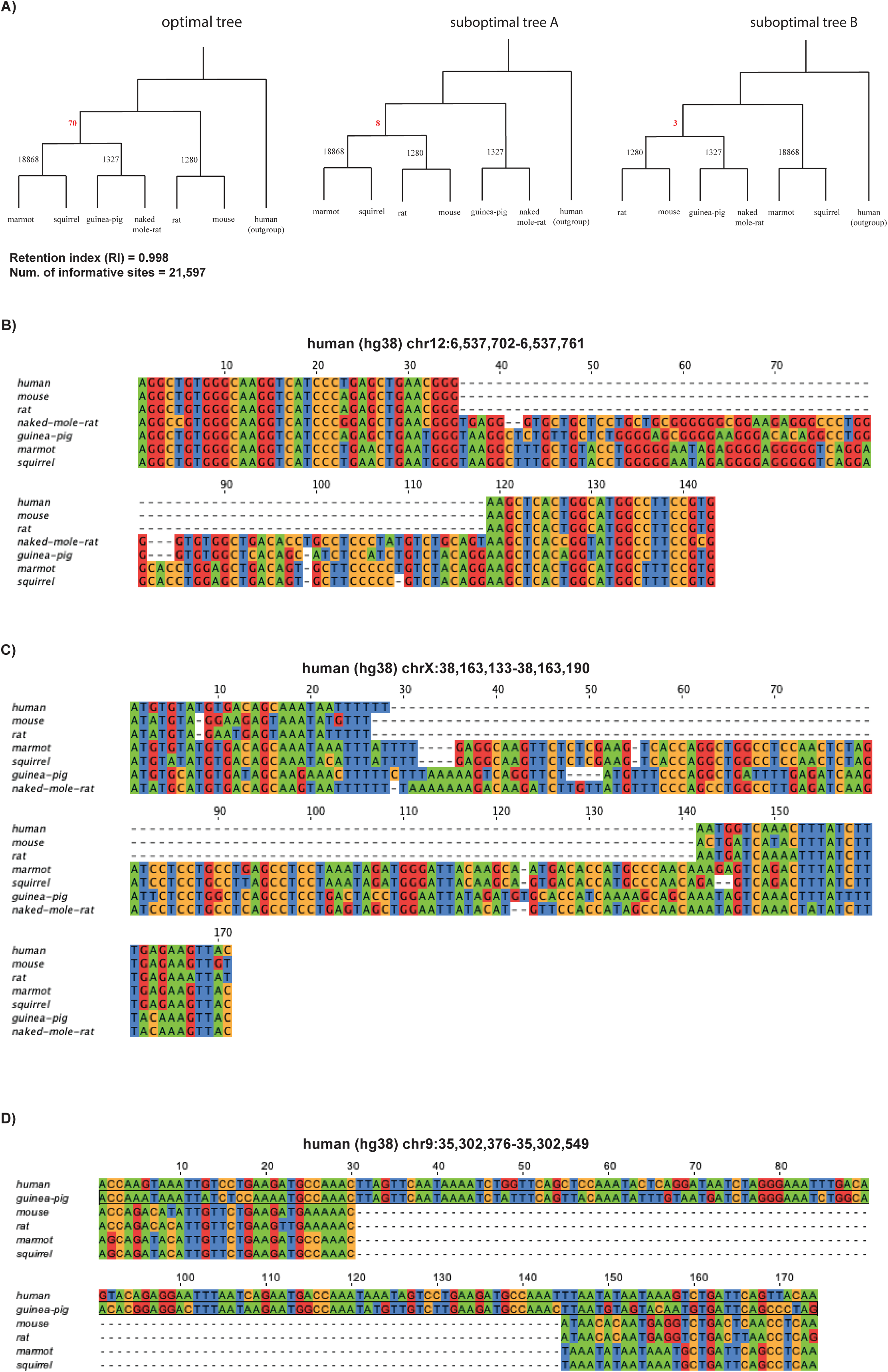
Champagne places Myomorpha basal to Sciuridae and Hystricomorpha. **A)** The maximum parsimony tree generated by PAUP* using Champagne’s character matrix for a subset of rodents (left), alongside two less parsimonious trees that reflect alternate branching relationships between Sciuridae, Myomorpha, and Hystricomorpha. Seventy indels support Myomorpha as basal., while only 8 and 3 support the other alternatives. **B)** A multiple sequence alignment for an 82bp insertion shared by mouse and rat. **C)** A multiple sequence alignment for a 107bp insertion shared by mouse and rat. **D)** A multiple sequence alignment for a 114bp deletion shared by mouse, rat, marmot, and squirrel. The character state of naked mole-rat was marked by Champagne as indeterminable (either because of an partially-gapped alignment in the deletion range, or because the tested sequence similarities did not meet Champagne’s minimum threshold). This supports panel A, suboptimal tree A.

## Discussion

A homoplasy-free character matrix has long been sought for phylogenetic studies to overcome the limitations of the current morphological and short sequence-based approaches, that contain large component of this non-phylogenetic signal. Previous efforts to find such a “perfect” character matrix have mostly relied on rare genomic events caused by transposable elements (TEs)^24, 26^. Such methods suffer from two limitations. First, the search for TE-based events has been very manual, with no efficient means developed of automation at the whole-genome scale. Second, events involving TEs, even though rare, are also suspected to suffer from a small level of homoplasy resulting from biological mechanisms^48^.

In this paper, our technique, Champagne, is concerned with *topologies* or *cladograms* rather than *trees*, since we exclude any molecular-clock-related inference from our analysis and focus on character matrix generation. Using the retention index (RI)^36^ on six sets of well-established topologies, we demonstrate that Champagne is largely homoplasy-free, with little or no non-phylogentic signal, which is in sharp contrast with both short sequence-based^39^ and morphological studies^43^, and overcomes a number of limitations of past approaches. First, by using pairwise whole-genome alignments to conservatively predict orthology of protein-coding genes, and further restricting the search to only intra-genic regions (which cover *>*35% of the human genome), Champagne performs genome-scale search, typically finding hundreds of large and rare genomic events, including, in large part, in the non-coding regions of the genome, where finding orthology is considered more challenging^49^. Second, Champagne is automated and easily scalable — Champagne requires gene annotation in a single known outgroup species and can work with unannotated genome assemblies for all target species. Champagne relies only on pairwise whole-genome alignments, which are much cheaper to compute than multiple-sequence alignments. In particular, for *N* ingroup species, Champagne requires only *N* pairwise alignments, one for each ingroup species paired with the outgroup. Using 9 primate species, we shows how Champagne can perform accurate, multi-species phylogenetic studies. Unlike methods involving only transposable elements, Champagne is oblivious to the biological mechanism or the sequence identities involved in its genomic events.

It is both theoretically expected and anecdotally shown (by the lack of current consensus) that some phylogenetic nodes are more difficult to resolve than others; as previously referenced, a considerable number of phylogenies have been either left unresolved or disputed. The ability of Champagne to produce a high-signal, low-noise (low-homoplasy) character matrix is necessarily constrained by the same biological phenomena that has historically made resolving such nodes difficult. The biological process that causes incongruence between gene trees and species trees will cause incongruence, or apparent homoplasy, in the character matrix produced by Champagne. The two primary biological processes that cause such incongruence are: incomplete lineage sorting (ILS), when rapid sequential speciation events prevent ancestral polymorphisms from being fully resolved into all resulting lineages^12^; and horizontal gene transfer^50^, when genetic information is transferred directly between different species. Champagne did find significant ILS involving large indels in the three Paenungulata species, which are believed to have undergone rapid speciation^46^. However, compared to Hobolth et al.^12^, who found ILS to be prevalent in *>*25% of the genome in the base-pair alignment of human, chimpanzee, and gorilla, Champagne observes zero indels that appear to result from ILS, while finding nearly 100 informative sites. We surmise that systematic biases in the character matrix of Hobolth et al.^12^ could have led them to overestimate genome-wide ILS in these species.

Most surprisingly, in this paper, we present a considerable set of indels that support the reevaluation of the relationship between Myomorpha, Hystricomorpha, and Sciuridae; our evidence suggests that Myomorpha should be considered basal to the latter two clades. Prior papers have presented alternate topologies, basing their conclusions upon a variety of evidence, including nuclear and mitochondrial DNA^38, 51^, morphological characters^43^, and SINEs^24^. Churakov et al.^24^ performed a SINE/indel screen of rodent genomic information, finding eight SINEs and six indels to support an early association of the Mouse-related and Guinea pig-related clades, with the Squirrel-related clade being the sister group. The authors note that “two SINE insertions and one diagnostic indel support an association of Hystricomorpha with the Squirrel-related clade”, suggesting that these conflicts might be explained by incomplete lineage sorting and hybridization. Champagne also searches for homoplasy-free indels but does so across 19,919 genes, resulting in a dataset that finds 70 indels in support of the positioning of Myomorpha as a sister group to Sciuridae and Hystricomorpha. Champagne, too, finds evidence supporting alternative topologies — 11 indels, in fact — and like Churakov et al, we believe that these are likely the result of ILS and potential hybridization. Given the lack of homoplasy inherent to its genome-wide derived characters, and 5 times more evidence, we argue that the Champagne character matrix is less prone to sampling bias than Churakov et al., and presents a compelling case to suggest that Myomorpha is, in fact, basal to Hystricomorpha, and Sciuridae.

Champagne is a highly general method that can easily be used on any sequenced set of species, along with an outgroup and its inferred gene set (derived even from gene-prediction or RNA-seq alone). Champagne promises to be much more homoplasy-free than morphological or single base-pair matrices. Moreover, while the ability to validate orthologous indels is expected to decay over large evolutionary distances, careful orthologous ancestral genomic region reconstruction^52^ promises to extend its reach even further back in time. A plethora of newly and soon-to-be sequenced species await analysis with Champagne.

## Methods

### Species set and gene set

In this study we used genome assemblies of 23 species (listed in Supplementary Table 1), and used Ensembl 86 (http://www.ensembl.org) for our reference (outgroup) species’ gene sets.

### Whole genome alignments and mapping orthologous genes

Once we selected a group of query (ingroup) species to study, we chose a known outgroup species for that group that also served as the reference. For each reference-query genome pair, Champagne used whole-genome pairwise alignments in the format of Jim Kent’s BLASTZ-based chains^33^ downloaded from the UCSC genome browser test server (https://hgdownload-test.gi.ucsc.edu/goldenPath/), or computed with the help of doBlastzChainNet utility (https://github.com/ENCODE-DCC/kentUtils) with default parameters for alignments not found on the server. For each reference gene, Champagne identified at most one orthologous chain in each query species, when it could do so with high confidence. First, it assigned every coding base in the canonical transcript of the reference gene to the highest-scoring chain (in terms of UCSC chain alignment scores) that overlaps with the base in its alignment. If the chain to which most bases were assigned was also the highest-scoring chain overlapping in its alignment by one or more base-pairs with the gene, then that chain was chosen as the best ortholog candidate, *C_b_* (see Figure 1, step 1). To ensure that there was no confusing paralog to *C_b_*, we required the UCSC alignment score of *C_b_* to be at least 20 times higher than any other chain overlapping with the gene by one or more base-pairs. To also ensure high synteny of *C_b_*, we required the number of bases in the aligning blocks of the chain *C_b_* be at least 20 times greater than the number of bases in the gene itself, i.e. *gene-in-synteny ≥* 20, where *gene-in-synteny* = length of *C_b_* / length of gene. We also required a unique *1-to-1 mapping* of coordinates between reference and query genomes, such that if two or more reference genes were mapped to the same query location, all overlapping mappings were discarded. If *C_b_* satisfied all above conditions, it was considered as the orthologous query chain containing the reference gene. In all remaining cases, no orthologous query chain was assigned for the reference gene.

### Identification and validation of insertions and deletions

Next, for each outgroup gene that mapped to a unique chain in more than one query species, Champagne scanned the query regions orthologous to the reference (outgroup) intragenic regions (exons as well as introns), moving through the outgroup-query chains simultaneously and identifying large (*>*75bp) indels from one-sided gaps in the chains. Specifically, a single-sided gap on the outgroup indicates either an insertion in query or a deletion in outgroup, while a single-sided gap on the query species indicates either a deletion in query or an insertion in outgroup (see Figure 1, step 2).

Upon finding an apparent indel in one such chain, Champagne located the corresponding coordinates in all other reference-query chains, and determined whether the indel event has occurred in the other query species by a combination of two methods: first, it confirmed the presence or absence of a similar-sized (within 10bp) single-sided gap in the other species; and second, it extracted species’ sequences within a fixed-size window range (of size *W* = 30bp) on either side of the indel and compared them directly (Supplementary Figure 1). For instance, if Champagne identified an insertion of size *δ* in query species *A* occurring at reference coordinate *X* (since a single-sided gap in the reference will start and end at the same coordinate), in order to verify the presence or absence of the insertion in another query species *B*, Champagne first checked that there is a single-sided gap of size *δ^′^*, where *|δ − δ^′^| ≤* 10, in the reference-query *B* chains at reference coordinate *X^′^*, within a 5bp margin from *X* (i.e. *|X − X^′^| ≤* 5bp). If such a gap was found, Champagne extracted the insertion sequence in both query *A* and *B*, and compared their sequence similarity. It also extracted a fixed-size ‘window’ sequence on either side of *X* and *X^′^* and compared them independently. If all of the sequence similarities exceeded our set threshold (determined as described below), Champagne assigned the indel a character state of ‘+’ (present) for species B, indicating that the insertion should be considered present. If the sequence similarities did not all exceed the threshold, Champagne assigned the indel a character state of ‘?’ (not confidently determinable). If no single-sided gap was found in species *B* near coordinate *X*, Champagne extracted species B’s window sequence on either side of *X* and compared it with species A’s window sequence; if the similarities both exceeded our threshold, the indel was assigned a state of ‘-’. Champagne also verified that the character state in the outgroup is actually the ancestral state (as opposed to an indel that has occurred independently in the outgroup) by requiring that at least one ingroup species aligns with high sequence similarity with the outgroup in the indel region and its surrounding windows without any large gaps. This verifies the ancestral state because we assume a very small probability of the independent occurrence of an indel at precisely the same locus in both the outgroup species and the ingroup species to which it aligns. Champagne discarded all sites where either the outgroup state could not be inferred to be the ancestral state, or where fewer than two query species had that indel.

For visual verification purposes (Figures 2-5), Champagne extracted the sequences of all species at the indel site and its surrounding windows, and used them to generate a multiple sequence alignment in the indel region using MUSCLE^53^.

### Dynamic threshold selection and evidence filtering

Recording the sequence similarity scores for each indel enabled the final step, in which Champagne tested a small range of minimum sequence similarity thresholds for insertions and deletions separately. We performed a parameter grid search over combinations of insertion and deletion thresholds in 0.25 intervals in the range [0.6, 0.7]. For each combination, we filtered out all indels that didn’t meet the stated thresholds across all species. Using the resulting evidence subsets, we then generated the most parsimonious topology using PAUP*, and calculated the ratio between the number of indels in support of alternate bifurcation hypotheses on internal nodes in that topology (per our definition of support outlined above). We optimized for the ratio between the number of indels that support the most- and second-most-supported bifurcation hypotheses on the ‘hardest’ node in the tree (the node with the lowest such ratio), selecting the thresholds that maximize this ratio. Crucially, we selected these thresholds regardless of what the optimal topology actually was.

### Topology inference and comparison baseline

Following this threshold selection step, Champagne filtered out all evidence that failed to meet the designated thresholds, and converted the labelled indels to a character matrix in NEXUS format (Figure 1, step 3), to infer the most parsimonious tree topology using PAUP*^28^ (Figure 1, step 4). To compare the retention indices of the topologies produced by Champagne with traditional approaches, we downloaded the single nucleotide sequence-based and morphology-based matrices (in NEXUS format) provided by Song et al.^39^ and O’Leary et al.^43^, respectively. From these matrices we extracted the rows corresponding to the same set of ingroup and outgroup species that were used by Champagne. We used PAUP* to generate the most parsimonious topology, specifying the outgroup species and using exhaustive search on each matrix, and recorded the associated retention index (RI) and the number of informative sites.

### Identifying evidence supporting a particular bifurcation

For each bifurcating branch in the tree, we also found the evidence in the Champagne matrix that supported the bifurcation. This was done as follows. For a branch which bifurcates into two sets of species, *A* and *B*, remaining ingroup species form another set *C*. An event was called supporting for this bifurcation if it indicated a shared insertion or deletion unique to species in *A* and *B*, not shared by any species in *C*. For shared insertions, we required at least one species in both *A* and *B* to be assigned a ‘+’, no species in either *A* or *B* to be assigned a ‘-’, at least one species in *C* to be assigned a ‘-’, no species in *C* to be assigned a ‘+’ and the outgroup to be assigned ‘-’. Similarly, for shared deletions, we required at least one species in both *A* and *B* to be assigned with a ‘-’, no species in either *A* or *B* to be assigned a ‘+’, at least one species in *C* to be assigned with a ‘+’, no species in *C* to be assigned a ‘-’ and the outgroup to be assigned ‘+’.

## Acknowledgements

We thank Hiram Clawson and the UCSC Genome Browser team for providing us the mammalian alignment chains. This work was funded by the NIH Grant R01HG008742, a Packard Foundation Fellowship, and a Microsoft Faculty Fellowship (to G.B.).

## SI Tables and Figures

**Supplementary Table 1.**
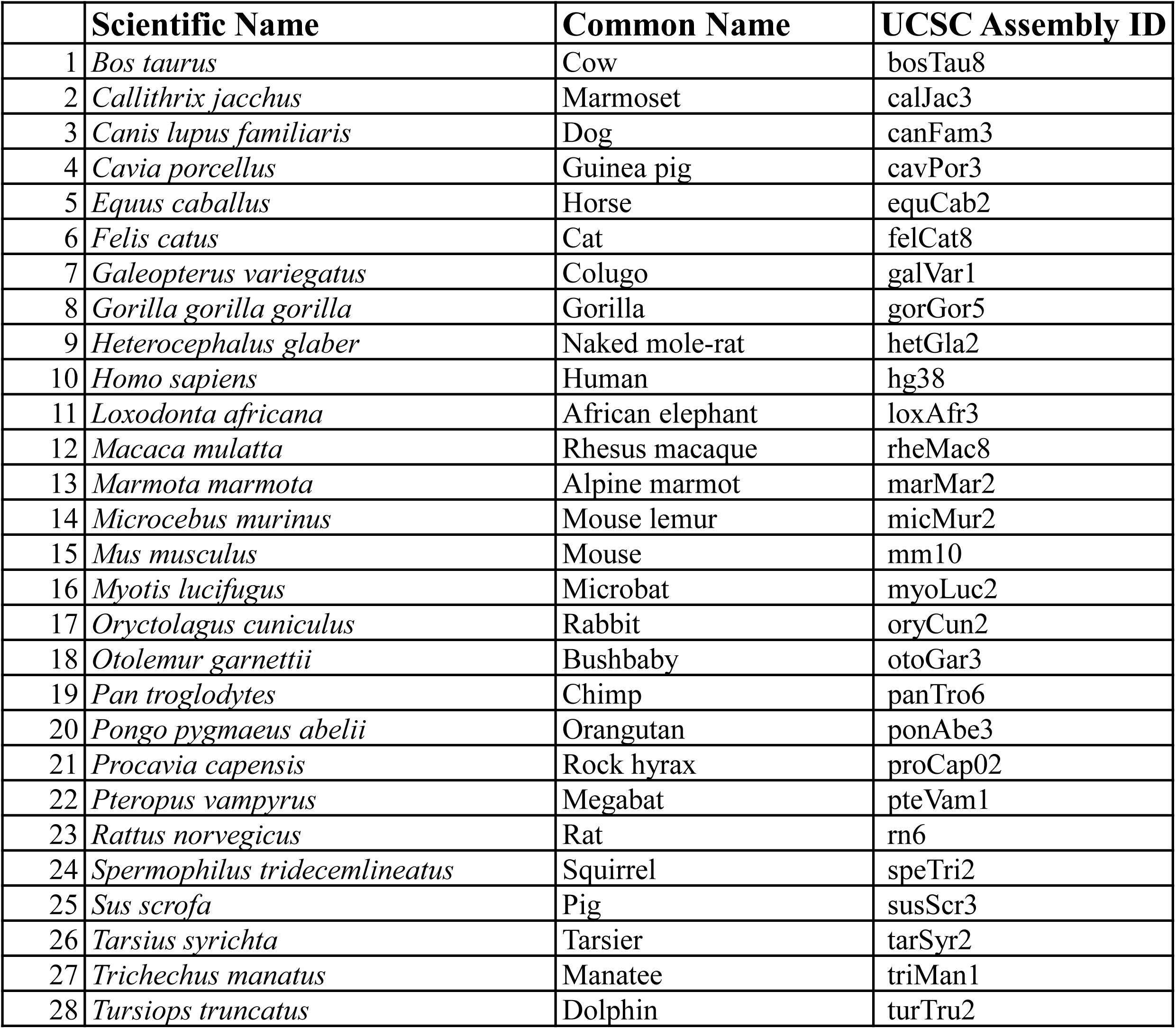
Scientific name, common name, and genome assembly name of all species used in this study.

|  | Scientific Name | Common Name | UCSC Assembly ID |
| --- | --- | --- | --- |
| 1 | *Bos taurus* | Cow | bosTau8 |
| 2 | *Callithrix jacchus* | Marmoset | calJac3 |
| 3 | *Canis lupus familiaris* | Dog | canFam3 |
| 4 | *Cavia porcellus* | Guinea pig | cavPor3 |
| 5 | *Equus caballus* | Horse | equCab2 |
| 6 | *Felis catus* | Cat | felCat8 |
| 7 | *Galeopterus variegatus* | Colugo | galVar1 |
| 8 | *Gorilla gorilla gorilla* | Gorilla | gorGor5 |
| 9 | *Heterocephalus glaber* | Naked mole-rat | hetGla2 |
| 10 | *Homo sapiens* | Human | hg38 |
| 11 | *Loxodonta africana* | African elephant | loxAfr3 |
| 12 | *Macaca mulatta* | Rhesus macaque | rheMac8 |
| 13 | *Marmota marmota* | Alpine marmot | marMar2 |
| 14 | *Microcebus murinus* | Mouse lemur | micMur2 |
| 15 | *Mus musculus* | Mouse | mm10 |
| 16 | *Myotis lucifugus* | Microbat | myoLuc2 |
| 17 | *Oryctolagus cuniculus* | Rabbit | oryCun2 |
| 18 | *Otolemur garnettii* | Bushbaby | otoGar3 |
| 19 | *Pan troglodytes* | Chimp | panTro6 |
| 20 | *Pongo pygmaeus abelii* | Orangutan | ponAbe3 |
| 21 | *Procavia capensis* | Rock hyrax | proCap02 |
| 22 | *Pteropus vampyrus* | Megabat | pteVam1 |
| 23 | *Rattus norvegicus* | Rat | rn6 |
| 24 | *Spermophilus tridecemlineatus* | Squirrel | speTri2 |
| 25 | *Sus scrofa* | Pig | susScr3 |
| 26 | *Tarsius syrichta* | Tarsier | tarSyr2 |
| 27 | *Trichechus manatus* | Manatee | triMan1 |
| 28 | *Tursiops truncatus* | Dolphin | turTru2 |

**Supplementary Figure 1.**
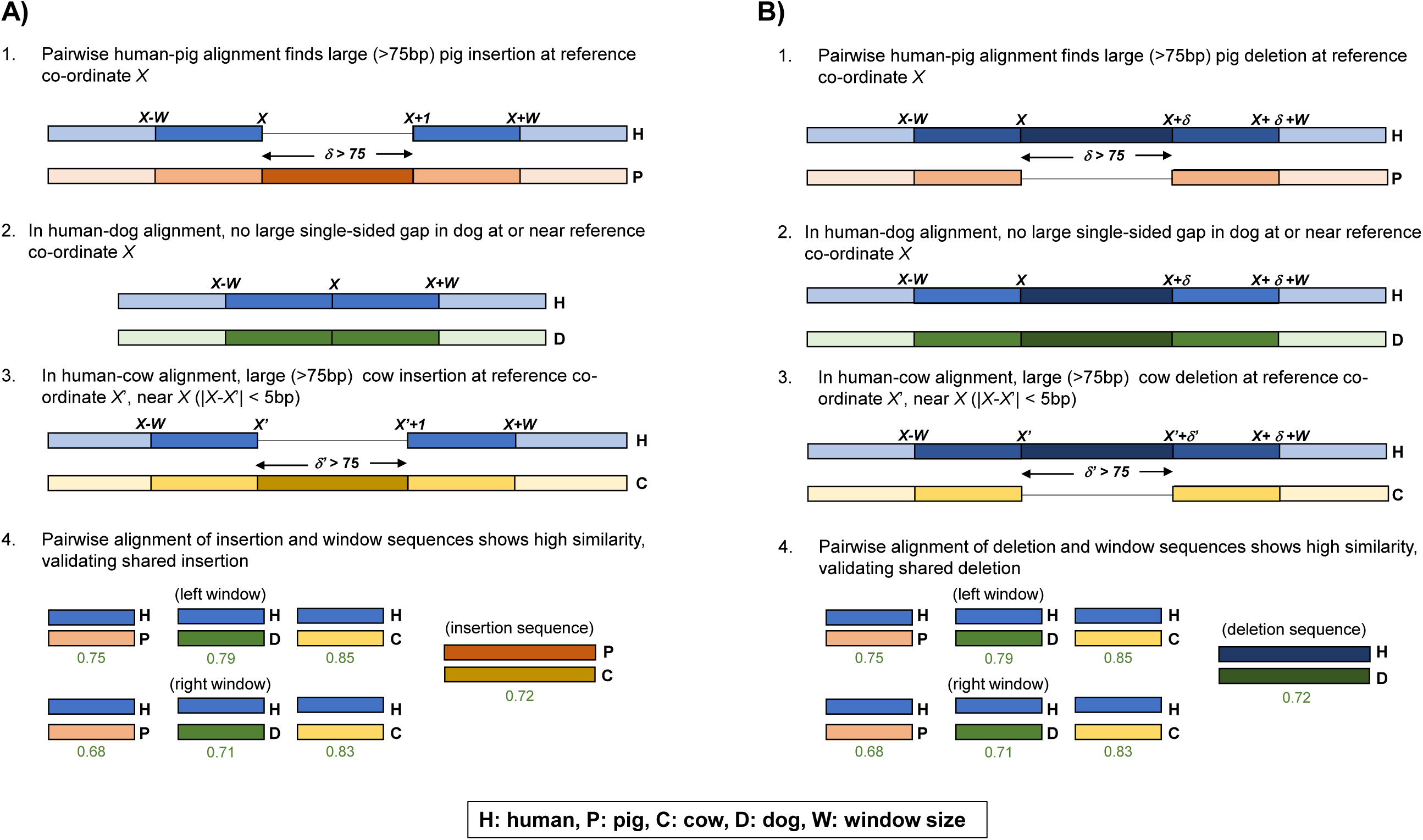
Champagne’s indel verification method. **A)** Shared insertion between pig and cow detected by Champagne that is absent in dog. (1) We first identify the presence of this insertion by finding a single-sided human gap in the human-pig orthologous chains, at human coordinate X. (2) Next, we find that there is no such single-sided gap in dog chain near X, we mark the insertion as likely absent in dog. (3) Next, we navigate to coordinate X in the human-cow chains, and check for a large (similar-sized) gap within at XI, within a 5bp range of X. Finding such a gap, indicating an insertion, we mark the insertion as likely present in cow. (4) Finally, we perform a direct sequence comparison for sequence similarity. We extract a W-sized ‘window’ (W = 30 in Champagne) sequence from either side of the insertion coordinate X in human, either side of the corresponding insertion coordinate in dog, and either side of the insertion itself in cow and pig. We also extract the sequence of the insertion itself in cow and pig. We then align the reference window sequences against each other species’ window sequences. Similarly, we align pig’s insertion sequence against cow’s insertion sequence. For each species in which we marked the indel as present, if the minimum sequence similarity for the left window, right window, and insertion (if the insertion is present) is greater than our stipulated threshold, we mark the species as definitively ‘+’. For each species in which we marked the indel as absent, if the sequence similarities for the left window and right window are greater than our stipulated threshold, we mark the species as definitively ‘-’. In either case, if a comparison fails the meet the threshold, we mark the species as ‘?’. **B)** Symmetrical process for finding shared deletions.

**Supplementary Figure 2.**
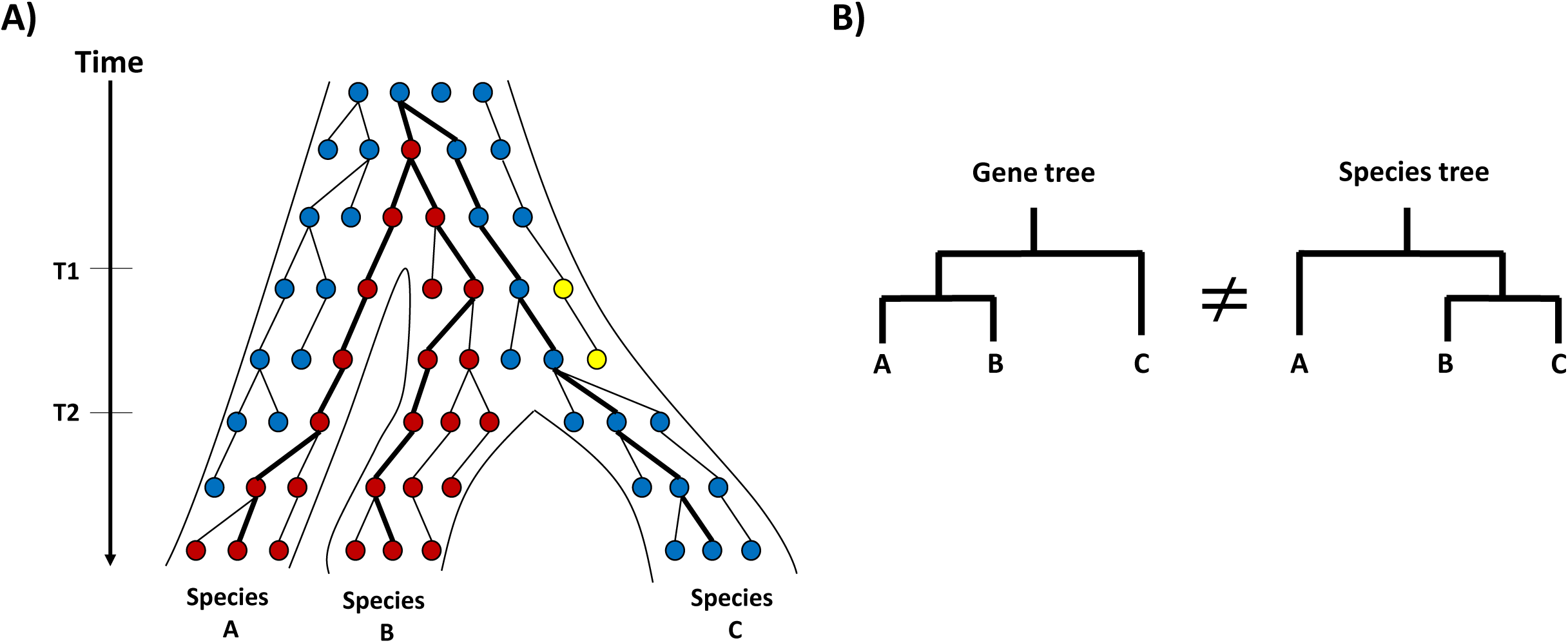
Gene tree incongruence caused by incomplete lineage sorting (ILS). **A)** Each dot represents the state (indicated by its color) of an allele at the same genomic site carried by a member of the population(s) evolving with time (vertical axis). Mutations (such as substitutions, insertions or deletions) cause new alleles (red and yellow) to appear in the population(s). Genetic drift can cause the allele frequency to increase (red) or decrease (yellow) in the population(s) over time. Two speciation events are shown to occur at times T 1 and T 2, resulting in three distinct species: A, B and C. **B)** The gene tree constructed from this allele (using the highlighted lineages in panel A) differs from the real species tree. This is due to incomplete lineage sorting, or the failure of species B, C that diverged on the right at time T 1 to “coalesce” the entire population to carry a single allele before another speciation occurs at T 2. The frequency of ILS in different alleles typically increases as the time between the two speciation events (T 1 and T 2) grows smaller.

